# Genetic basis of thermal plasticity variation in *Drosophila melanogaster* body size

**DOI:** 10.1101/268201

**Authors:** E Lafuente, D Duneau, P Beldade

## Abstract

Body size is a quantitative trait that is closely associated to fitness and under the control of both genetic and environmental factors. While developmental plasticity for this and other traits is heritable and under selection, little is known about the genetic basis for variation in plasticity that can provide the raw material for its evolution. We quantified genetic variation for body size plasticity in *Drosophila melanogaster* by measuring thorax and abdomen length of females reared at two temperatures from a panel representing naturally segregating alleles, the Drosophila Genetic Reference Panel (DGRP). We found variation between genotypes for the levels and direction of thermal plasticity in size of both body parts. We then used a Genome-Wide Association Study (GWAS) approach to unravel the genetic basis of inter-genotype variation in body size plasticity, and used different approaches to validate selected QTLs and to explore potential pleiotropic effects. We found mostly “private QTLs”, with little overlap between the candidate loci underlying variation in plasticity for thorax versus abdomen size, for different properties of the plastic response, and for size versus size plasticity. We also found that the putative functions of plasticity QTLs were diverse and that alleles for higher plasticity were found at lower frequencies in the target population. Importantly, a number of our plasticity QTLs have been targets of selection in other populations. Our data sheds light onto the genetic basis of inter-genotype variation in size plasticity that is necessary for its evolution.

**Significance Statement:** The environmental conditions under which development takes place can affect developmental outcomes and lead to the production of phenotypes adjusted to the environment adults will live in. This developmental plasticity, which can help organisms cope with environmental heterogeneity, is heritable and under selection. Plasticity can itself evolve, a process that will be partly dependent on the available genetic variation for this trait. Using a wild-derived *D. melanogaster* panel, we identified DNA sequence variants associated to variation in thermal plasticity for body size. We found that these variants correspond to a diverse set of gene functions. Furthermore, their effects differ between body parts and properties of the thermal response, which can, therefore, evolve independently. Our results shed new light onto a number of key questions about the long discussed genes for plasticity.

## INTRODUCTION

Body size has a great impact on the performance of individuals (1, 2), as well as that of species (3). Diversity in this trait is shaped by the reciprocal interactions between the developmental processes that regulate growth, and the evolutionary forces that determine which phenotypes increase in frequency across generations (4). Body size varies greatly within and between populations (2, 5) and is controlled by both genetic and environmental factors (6–9). Studies in different animal models have provided insight about the selection agents that shape the evolution of body size (e.g. predators (10, 11), mates (12), thermal regimes (13, 14)), and about the molecular mechanisms that regulate body size and body proportions during development (15–18).

Body size is also a prime example of the environmental regulation of development, or developmental plasticity (19, 20), and it is influenced by different factors, including nutrition and temperature. This plasticity can help organisms cope with environmental heterogeneity and, as such, can have major implications for population persistence and adaptation (20–22). Thermal plasticity in body size is ubiquitous among insects (23–25), with development under colder temperatures typically resulting in larger bodies, which is presumably advantageous for thermalregulation (1, 26). The environmental dependency of body size, and other plastic traits, is often studied using reaction norms, in which phenotypic variation is plotted as a function of gradation in the environment (27). The properties of these reaction norms, including their shapes and slopes, can differ between genotypes (28–30), and the genes underlying such variation can fuel the evolution of plasticity. Little is known about what these genetic variants are and what types of functions they perform (e.g. perception of environmental cues, conveying external information to developing tissues, or executing actions in developing plastic organs). It is also unclear to what extent the loci contributing to variation in thermal plasticity in size are the same for different body parts, and whether the loci contributing to variation in size plasticity are the same that underlie inter-individual variation in body size at a given temperature.

Studies in *D. melanogaster* have provided much insight about the evolution and development of body size, body proportions, and body size plasticity (7, 31–38). Size differences among populations, including clinal (31, 39) and seasonal variation (40), and among individuals within a population, are due to the effects of genes, environment, as well as genotype-by-environment interactions (33, 41–43). While we have increasing detailed knowledge about the genetic basis of adaptation and of natural variation for many adaptive traits in *D. melanogaster* and other species (6, 43–45), little is known about the genetic basis of variation in plasticity. Widely-accessible mapping panels (46, 47) allowed the dissection of the genetic architecture of various quantitative traits in *D. melanogaster* (48–50), including body size (51). However, with a few recent exceptions (52–55), the genetic basis of phenotypic variation has been investigated under a single environmental condition, precluding assessment of differences between environments and of the genetic basis of plasticity itself. Series of isogenic lines from these mapping panels can be reared under different conditions to characterize reaction norms and ask about the genes that harbor allelic variation for their properties (53).

Here, we use a panel of *D. melanogaster* lines representing naturally segregating alleles from one natural population, the DGRP (45, 47), to characterize genetic variation for thermal plasticity in thorax and abdomen size, and to identify loci contributing to variation in the slopes of their thermal reaction norms. We document correlations between body size and body size plasticity, as well as correlations with other traits, using published data for the same panel. We also ask about the extent of overlap between QTLs for size and for size plasticity, and between QTLs for size plasticity of the different body parts. We then use different approaches to validate and further characterize the role of selected QTLs, and to ascertain their pleiotropic effects.

## RESULTS

We measured thorax and abdomen length in adult females from different genotypes reared at two temperatures. We quantified effects of genotype (G), environment (E), and genotype-by-environment (GxE) interactions on body size (Fig. 1), and explored correlations between body parts and between temperatures (Fig. 2). We then used a GWAS approach to identify DNA sequence polymorphisms associated with variation in body size plasticity (Fig. 3). Ensuing functional analyses of candidate QTLs validated and clarified their role in body size variation at different temperatures (Fig. 4).

**Fig. 1.**
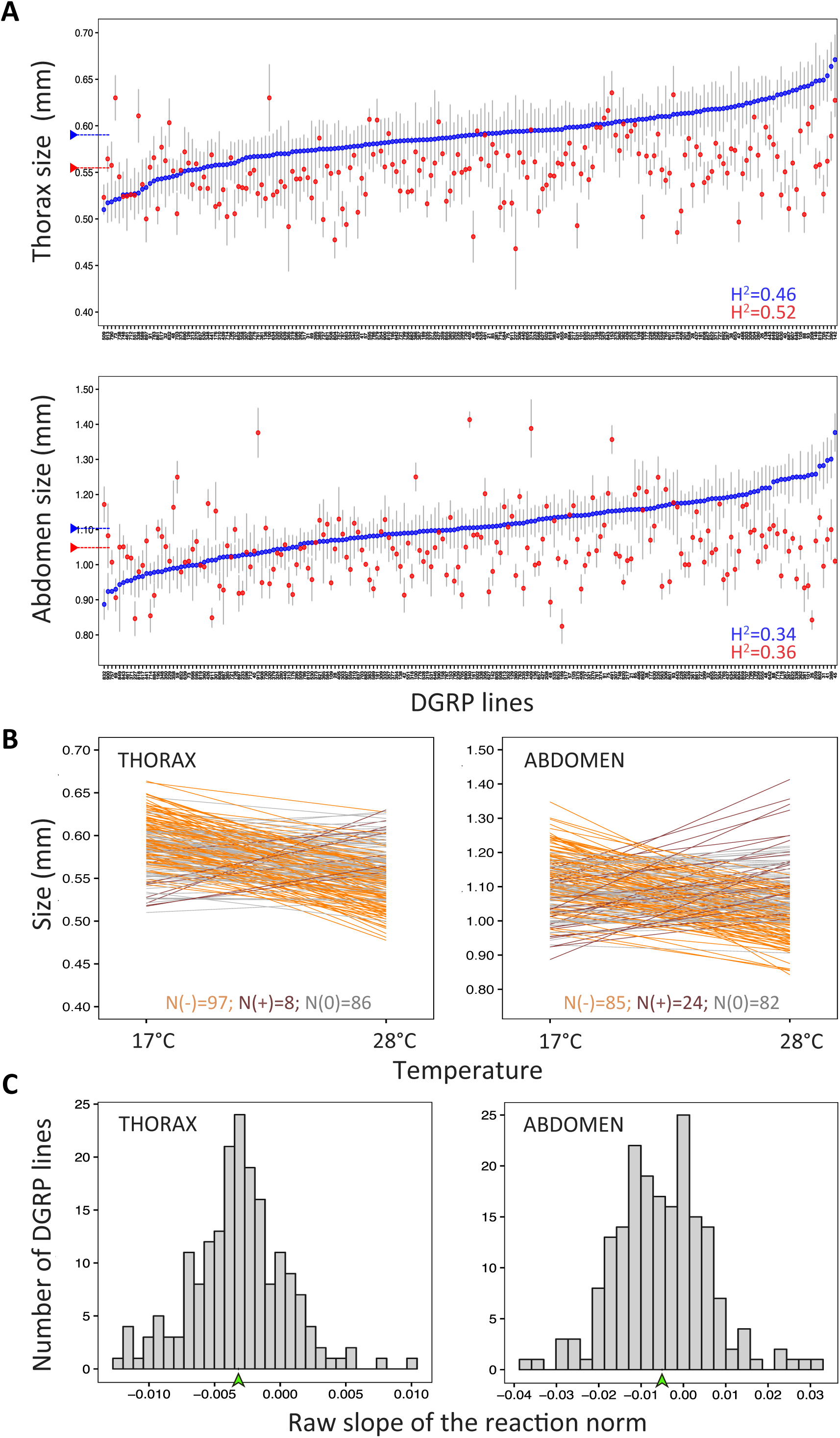
Natural genetic variation in size and size plasticity. **A**. Means and 95% confidence intervals (Y axis) for thorax (upper panel) and abdomen (lower panel) size in the DGRP lines (X axis) reared either at 17°C (blue) or at 28°C (red). DGRP lines are ranked by their mean size at 17°C. Dashed horizontal bar represents the mean value for all DGRP lines at a given temperature. Mean values (*μ)* and broad sense heritability (H^2^) estimates per body part and temperature can be found in Table 1. **B**. Reaction norms for thorax and abdomen sizes (Y axis) across temperatures (X axis) plotted as the regression fit for the model lm (*Size ~ Temperature*) for each DGRP line. Reaction norms are colored by in relation to slope: slopes significantly different from zero are orange when positive and brown when negative, while slopes that were not significantly different from zero are gray (alpha=0.05). Counts of each are shown on the bottom of each graph. Broad sense heritability estimates were: *H*^*2*^=0.33 (thorax plasticity) and *H*^*2*^=0.49 (abdomen plasticity). **C**. Frequency distribution for the raw value of the slope of the reaction norm in the DGRP lines. The mean value for the raw slope of all DGRP reaction norms is indicated with a green arrowhead.

**Fig. 2.**
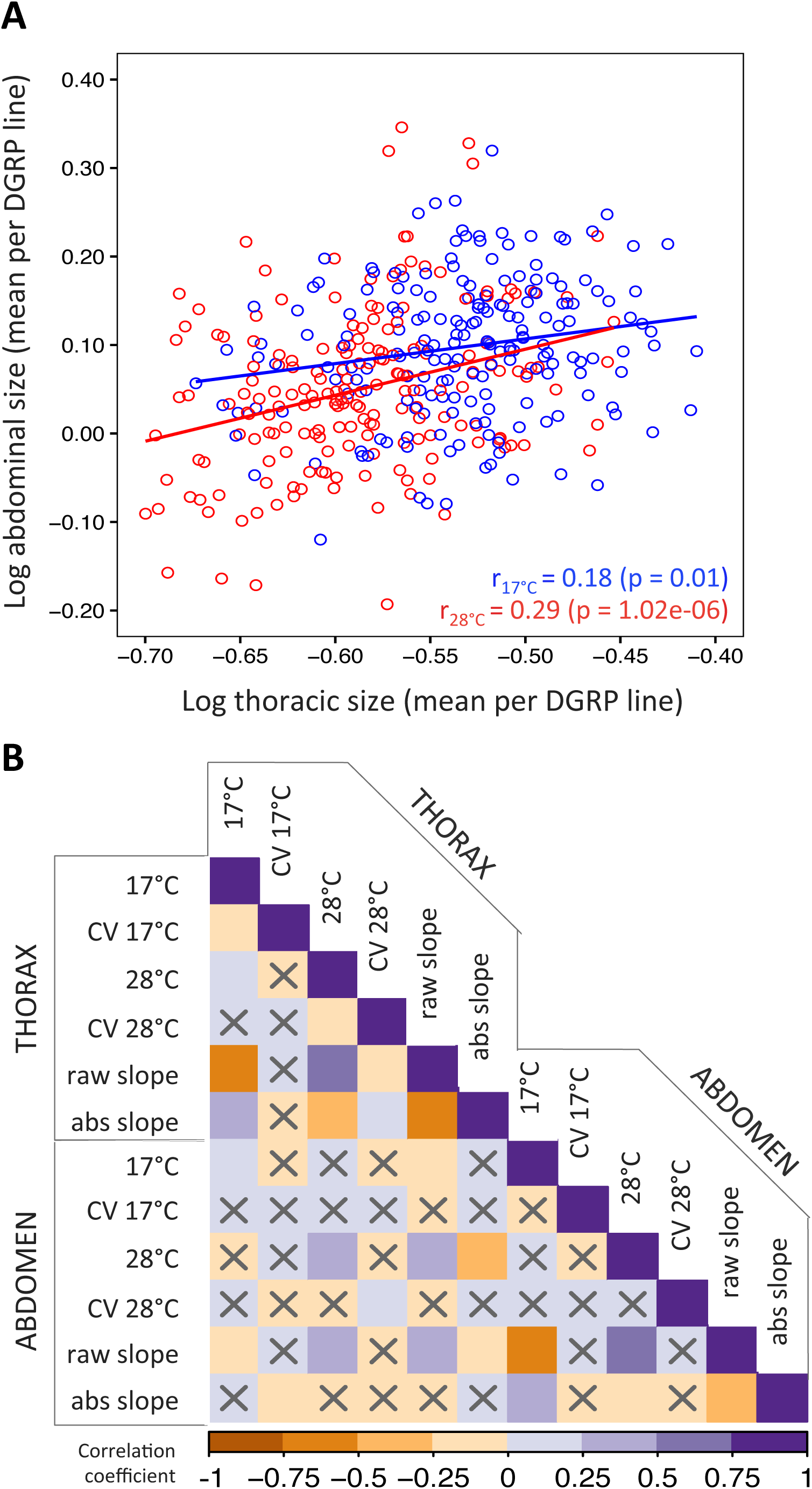
Phenotypic co-variation in size and size plasticity. **A**. Thoracic (X axis) and abdominal (Y axis) mean size per DGRP line at 17°C (blue) and at 28°C (red). Pearson correlation, r=0.18 (p-value=0.01) and partial Pearson correlation, r=0.34 (p-value= 1.02e-06) for 17°C, Pearson correlation r=0.29 (p-value<2e-16) and partial Pearson correlation r=0.34 (p-value<2e-16) for 28°C. **B**. Heat map of Pearson’s correlation coefficients between our measurements in thoraxes and abdomens: mean sizes and coefficients of variation (CV) at each temperature and raw and absolute slopes of the reactions norms. Non-significant correlations (p-value < 0.01) are indicated with an ‘X’.

**Fig. 3.**
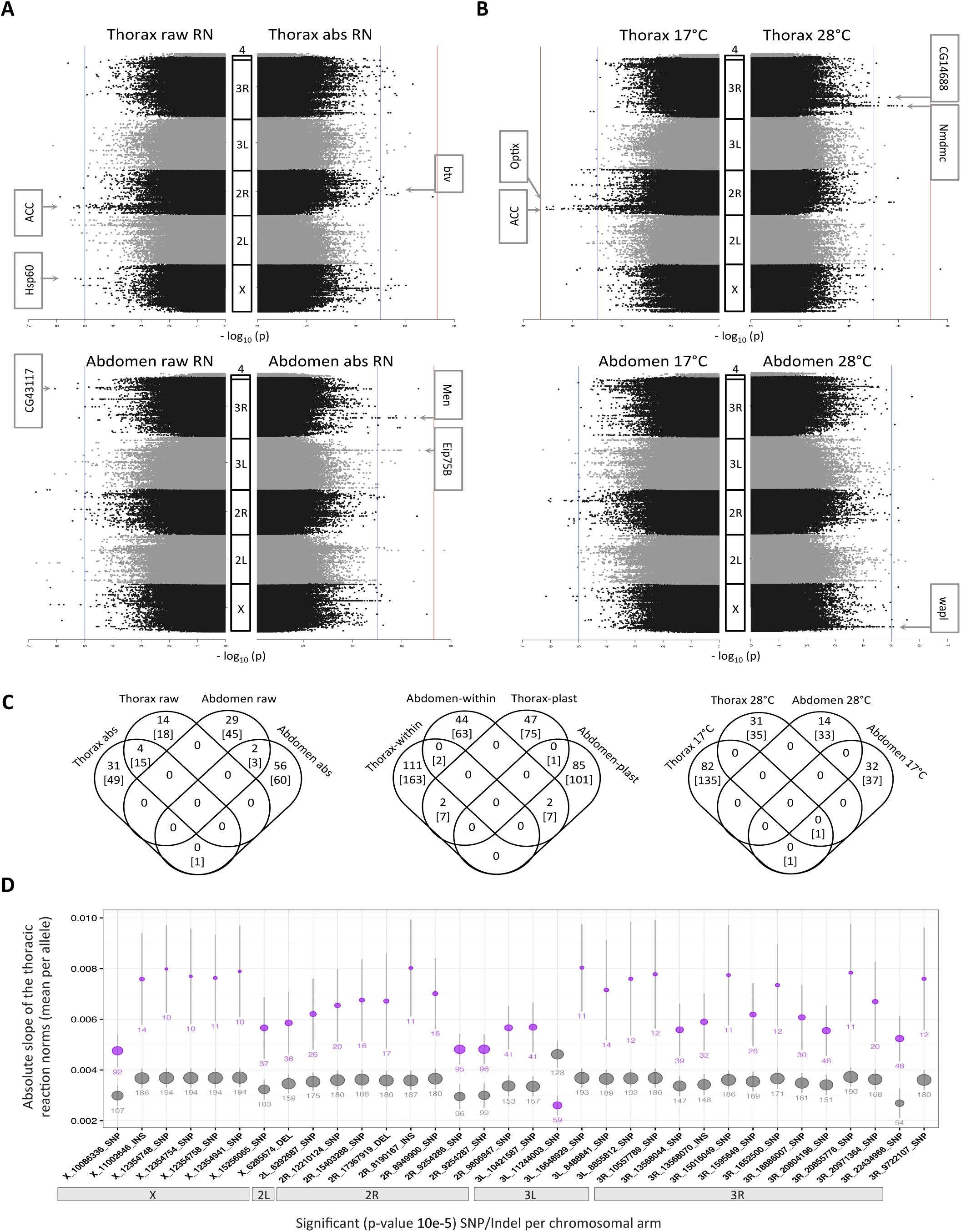
Genetic variants influencing size and size plasticity. **A-B**. Manhattan plots corresponding to the eight GWAS analyses performed. Horizontal lines are p-value<10e-5 (blue) and p-value<10e-8 (red). Gene names are shown for a subsample of significant SNPs/Indels, which were selected based on p-value, putative variant effect and associated genes (Table S2 and S3). **A**. GWAS for variation in size plasticity in thoraxes (upper panels) and abdomens (lower panels) and for either the raw (left) or the absolute (right) slopes of the reaction norms. **B**. GWAS for variation in size in thoraxes (upper panels) and abdomens (lower panels) at 17°C (left panels) or 28°C (right panels). **C**. Venn diagrams showing the number of candidate SNPs/Indels (number outside the brackets) and candidate genes (number within brackets) harboring those polymorphisms identified in the different GWAS. **D**. Mean and 95% confidence interval of the absolute slope of the reaction norms for thorax size (Y axis) per allele (major allele in gray, minor allele in magenta) for each candidate plasticity QTL along the chromosomal arms (X axis). The position and identity of the polymorphisms in this figure is given by their annotation with Genome Release v.5.

**Fig. 4.**
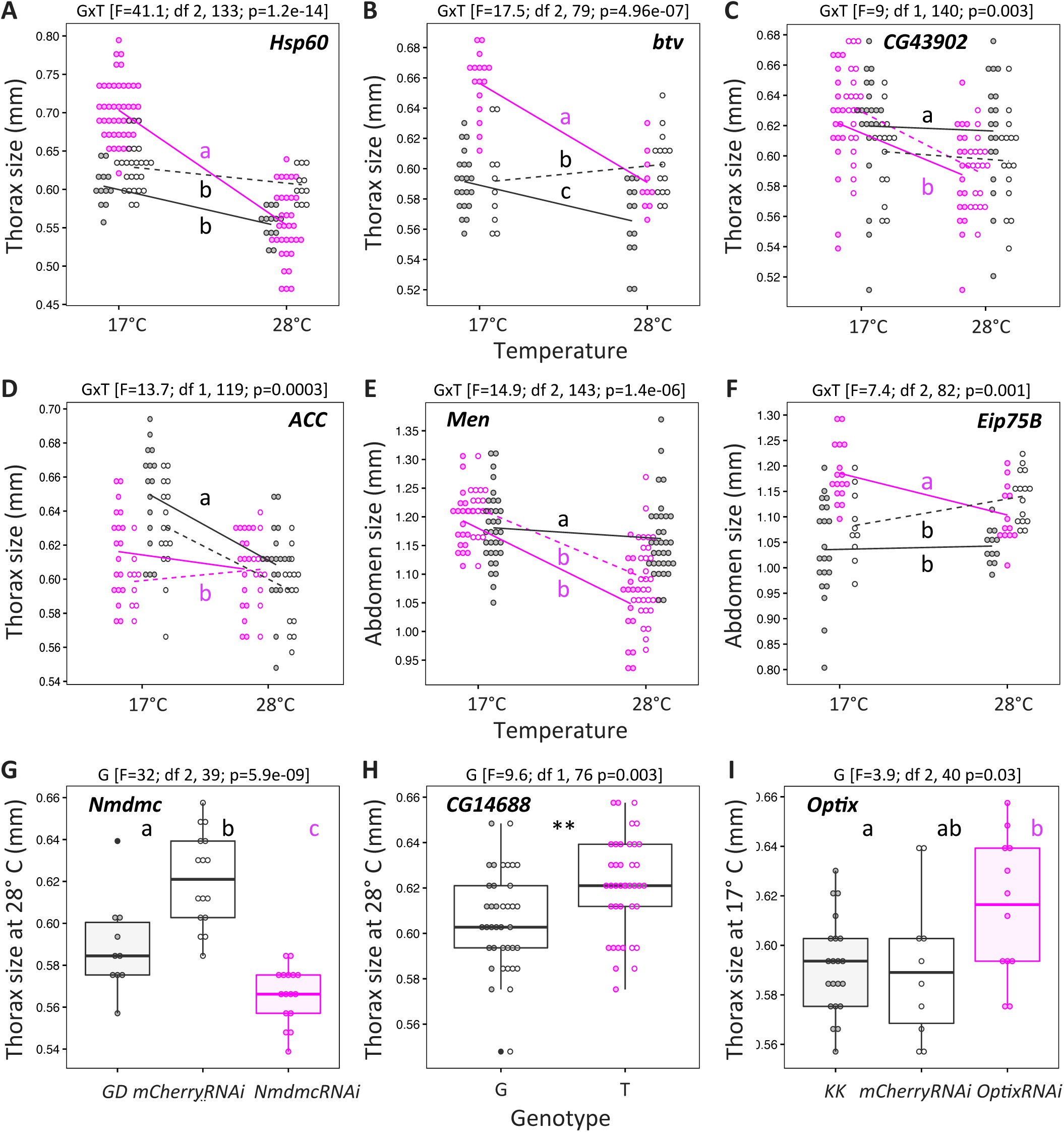
Functional validation of selected QTLs from GWAS analyses. For validations via mutant or RNAi, genotypes with impaired gene function are shown in magenta and control genotypes are shown in black. Similarly, for validations via Mendelian Randomization (MR), the two populations fixed for the minor allele are shown in magenta and the two populations for the major allele are shown in black. **A**. Reaction norms in mutant *Hsp60A/+* and controls Canton-S (filled circles, solid line) and *Fm7a/Canton-*S (empty circles, dashed line). **B**. Reaction norms in *btv-RNAi/bab-Gal4* and control lines *KK* (filled circles, solid line) and *mCherry-RNAi/bab-Gal4* (empty circles, dashed line). **C**. Reaction norms in the four MR populations for SNP X:10192303 within gene *CG43902*. **D**. Reaction norms in the four MR populations for SNP 2R:7983239 within gene *ACC*. **E**. Reaction norms in *Men* mutants *MenEx3/+* (filled circles, solid line) and *MenEx55/+* (empty circles, dashed line) and control line *w1118*. **F**. Reaction norms in *Eip75B-RNAi/bab-Gal4* and control lines *KK* (filled circles, solid line) and *mCherry-RNAi/bab-Gal4* (empty circles, dashed line). **G**. Size at 28°C in *Nmdmc-RNAi/tub-Gal4* and control lines GD (filled circles) and *mCherry-RNAi/tub-Gal4* (empty circles). **H**. Size at 28°C in the four MR populations for SNP 3R:10678848 within gene *CG14688*. **I**. Size at 17°C in *Optix-RNAi/bab-Gal4* and control lines KK (filled circles) and *mCherry-RNAi/bab-Gal4* (filled circles). For the validations of plasticity QTLs, we tested the model lm (*Size ~ Genotype*Temperature)* and for the validations of within-environment QTLs we tested the model lm (*Trait ~ Genotype*). Results from the statistical models are shown above each plot and, when significant, indicated by asterisks in the plot (where p-values < 0.001 and < 0.01 are denoted by ‘***’ and ‘**’, respectively). Differences for more than two groups were estimated by post hoc comparisons (Tukey’s honest significant differences) and are indicated by different letters in each plot (p-value<0.01). For all tested SNPs/genes, the phenotype of the DGRP lines carrying the minor versus the major allele at the target QTL can be found in Fig. S7.

### Partitioning variation in body size

To assess the contribution of genotype and temperature to body size variation, we quantified length of abdomens and thoraxes (Fig. 1A and S1A) of adult females from ~196 isogenic lines reared at either 17°C or 28°C (Dataset 1, Table S1). We found significant differences between DGRP genotypes and between developmental temperatures, as well as significant GxE interaction effects (thermal plasticity) for the size of both body parts (Fig. 1A and S1C). We also found variation between individuals of (presumably) the same genotype and same rearing temperature; the coefficients of variation varied between 0.6 and 15 for thorax measurements and between 0.3 and 23.8 for abdomen measurements (Table 1). Broad-sense heritabilities for body size (at 17°C and 28°C) and body size plasticity (between-environment variation) varied between 30 and 50% (Table 1).

**Table 1.** Summary measurements and broad-sense heritability estimates for size and size plasticity ^1^ Body part (T=thorax, A=abdomen), ^2^ subsets used to perform calculations: temperature for within-environment parameters and raw slope of the reaction norm for plasticity parameters, ^3^ minimum and maximum size mean value (in mm) per group, ^4^ minimum and maximum coefficient of variation per group, ^5^ variance associated to genetic differences, ^6^ variance associated to residual differences, ^7^ variance associated to genotype-by-temperature interaction component, ^8^ total variance, ^9^ heritability estimates. See materials and methods for details on the performed calculations.

| Body part <sup>1</sup> | Trait <sup>2</sup> | Mean <sup>3</sup> | CV <sup>4</sup> | Genetic variance <sup>5</sup> | Residual variance <sup>6</sup> | GxT variance <sup>7</sup> | Total variance <sup>8</sup> | Heritability <sup>9</sup> |
| --- | --- | --- | --- | --- | --- | --- | --- | --- |
| T | 17°C | 0.51-0.67 | 0.6-12.1 | 7.98e-04 | 1.6e-03 | NA | NA | 0.34 |
|  | 28°C | 0.47-0.64 | 0.9-15 | 8.7e-04 | 1.6e-03 |  |  | 0.36 |
| A | 17°C | 0.89-1.38 | 0.4-17.7 | 6.6e-03 | 7.6e-03 |  |  | 0.46 |
|  | 28°C | 0.82-1.41 | 0.3-23.8 | 8.7e-03 | 7.9e-03 |  |  | 0.52 |
| T | slope | -3.18e-03 | NA | 2.0e-04 | 1.6e-03 | 6.0e-04 | 2.3e-03 | 0.33 |
| A | slope | -5.05e-03 |  | 12.3 | 92.5 | 75.2 | 180.1 | 0.49 |

We computed a correlation matrix to assess relationships between the different components of variation in body size (Fig. 2). We found a significant positive correlation between thorax and abdomen size within each rearing temperature (Fig. 2A, B). We also found significant positive correlations between our measurements of thorax (both temperatures), but not abdomen size, and the size of several body parts measured in other studies (51) for the same genotypes reared at 25°C (Fig. S2A). To address the question of association between body size and fitness, we measured correlations between our measurements of body size and a series of fitness related traits quantified in other studies of the DGRPs (46, 49, 50, 56). We found significant correlations with chill coma recovery (thorax size at both temperatures and abdomen size at 28°C) and with survival upon infection with *Metarhizium anisopliae* fungi (abdomen size at 17°C), but not with longevity, starvation resistance, and other immune-defense traits (Fig. S2B). To address the question of what might explain the extent of inter-individual variation within genotype and temperature, we calculated correlations between our measurements of CV and of size. We found the CV to be: 1) positively correlated between body parts for flies reared at 28°C, but not for those reared at 17°C, and 2) negatively correlated with mean size for thoraxes, but not for abdomens (Fig. 2B).

### Thermal reaction norms for body size

Using reaction norms, we studied the extent and properties of thermal plasticity for body size in the DGRP lines (Fig. 1B). We calculated the slope of the regression lines for size across temperatures for each body part and DGRP genotype, and found genetic variation for both the intercept and slope of the reaction norms (Fig. 1B, Table S1). From each reaction norm, we extracted two properties of the thermal plasticity in body size: i) the absolute value of the slope, describing only the magnitude of the response to temperature, and ii) the raw value of the slope, which describes magnitude and direction of the response (Fig. 1C and S1D). Using the reaction norms for 191 DGRP lines, we identified plastic and non-plastic genotypes. Slopes of the thermal reaction norm were significantly different from zero for 55% and 57% of the lines for the size of thoraxes and abdomens, respectively, with most of the plastic genotypes having smaller sizes when reared at higher temperatures (Fig. 1B). However, we also found plasticity in the opposite direction (i.e. genotypes with smaller flies at lower temperatures), corresponding to a positive significant slope for the thermal reaction norms: 8% of the DGRP reaction norms for the thorax and 22% for the abdomen (Fig. 1B). Even though the levels of thermal plasticity for thorax and abdomen size were significantly positively correlated (Fig. 2B), lines having the highest levels of plasticity for one body part were not necessarily the most plastic for the other body part (Fig. S1B). Furthermore, for thorax, but not abdomen, we found that genotypes with larger CVs had steeper negative reaction norms. Finally, we also found no correlation between our size plasticity measurements and various fitness-related traits measured in the DGRPs (Fig. S2B), including longevity (49), starvation resistance, chill coma recovery (46), and immune-defense traits (50, 56).

### Genetic basis of variation in body size plasticity

We used a GWAS approach to identify DNA sequence polymorphisms associated with variation in thermal plasticity for thorax and abdomen size in the DGRP. Because the loci carrying allelic variation for the direction and extent of environmental responsiveness are not necessarily the same, we used both the raw and absolute values of the slopes of the DGRP reaction norms as quantitative traits (Fig. 3A and S3). The candidate QTLs significantly associated with variation in plasticity were typically only so in relation to a single property of the reaction norm (raw or absolute slope) or body part (thorax or abdomen; Fig. 3C, Table S2). We also found that these allelic variants fell within different genomic regions (e.g. UTR, intronic, coding) within or nearby 192 different putative genes (Tables 2 and S2). Gene ontology enrichment analysis of the candidate QTLs showed an over-representation of vesicle-mediated processes (e.g. phagocytosis and endocytosis; Fig. S6A), while network enrichment analyses (protein-protein interaction network followed by a KEGG pathways enrichment analysis) revealed an over-representation for SNARE interactions and Notch pathways (Fig. S6B), both of which have been implicated in diverse biological functions (57–60). We also found that in the vast majority of cases, alleles associated to increased environmental responsiveness were at lower frequencies in the DGRP (Fig. 3D and S5).

**Table 2.**
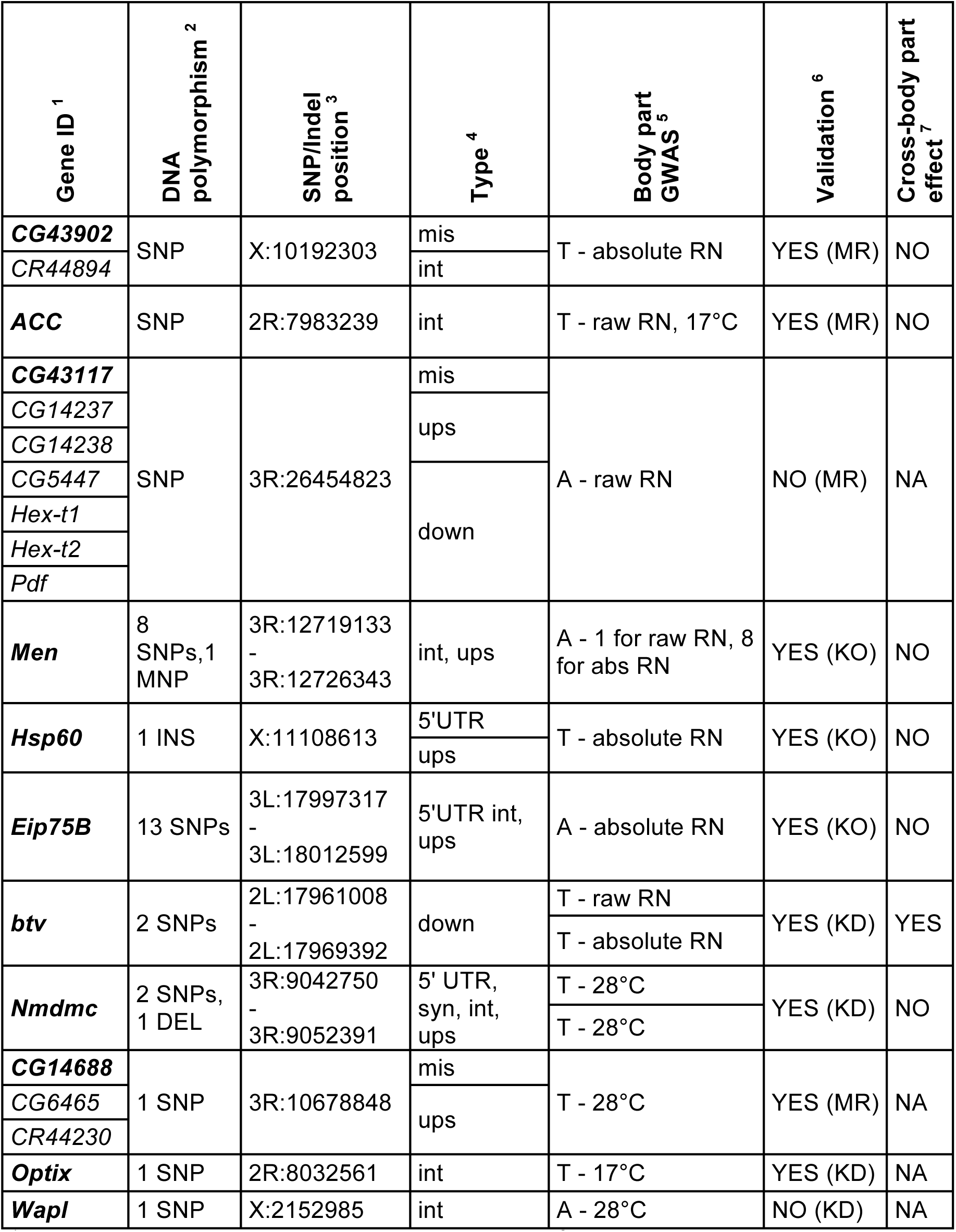
Details of candidate QTLs selected for functional validation ^1^ Putatively affected gene given as ID gene symbol, ^2^ type of polymorphism (SNP=Single Nucleotide polymorphism, INS=Insertion, DEL=Deletion, MNP= Multiple Nucleotide Polymorphism), ^3^ genomic position (FlyBase release FB2017_06 (98)), ^4^ putatively affected gene regions (*mis*=coding missense substitution, *int*=intronic, *ups*=upstream, *down*=dowstream, *syn*=coding synonymous substitution), ^5^ body part and trait for which the QTL was a hit (T=thorax, A=abdomen), ^6^ results of the validation (“*YES/NO”* for whether validation confirmed or not the role the target QTL on phenotype; results in Fig. 4 and S7) with the approach used (*MR*=Mendelian Randomization, *KO*=mutant, *KD*=RNAi), ^7^ results of the test for pleiotropic effects (“*YES/NO*” for whether we found cross body part effects; results in Fig S7).

To explore to what extent loci that contribute to variation in size plasticity also contribute to variation in body size within environments, we also performed GWAS analyses using body size at 17°C and 28°C as quantitative traits (Fig. 3B and S4, Table S3). This analysis revealed QTLs that were mostly environment- and body part-specific (Fig. 3C, Table S3), including no overlap between our candidate QTLs associated with variation in thorax and abdomen size and those reportedly associated with head size at 25°C (51). Moreover, we found little overlap between candidate QTLs contributing to variation in size plasticity and those contributing to within-environment size variation (Fig. 3C), but some overlap in terms of network enrichment (Fig. S6). None of our size traits or plasticity therein was affected by chromosomal inversions (p-value > 0.01), or by the genetic relatedness among DGRP lines (low and non-significant coefficients of phylogenetic signal Blomberg’s K and Pagel’s λ; Fig. S2C).

### Validation of selected GWAS hits

We selected a number of significant QTLs for validation via different approaches (Fig. 4, Table 1, Dataset 2). To test selected candidate genes, we used available null mutants and inducible gene knock-downs (with RNA interference using the Gal4/UAS system). If the candidate gene affects plasticity, we expected to see a difference in thermal reaction norms between control genotypes and those with abolished or reduced candidate gene function. To test specific significant SNPs/Indels, we used a SNP-based validation approach, hereafter called Mendelian Randomization (see Materials and Methods), that allowed comparisons between genotypes fixed for the candidate SNP (minor versus major alleles) but not for any other significant SNP for the same trait. If the candidate SNP affects plasticity, we expected a difference in slope of the reaction norms between newly-established genotypes carrying the minor versus the major allele at the target SNP.

Using these methods, we confirmed a role in thermal plasticity for six out of seven candidate QTLs (Fig. 4 and S7, Table 2). For genes *Hsp60* (Fig. 4A), *btv* (Fig. 4B), *Men* (Fig. 4E), and *Eip75B* (Fig. 4F), plasticity was different between genotypes with impaired gene function (knock-out or knock-down) versus controls. For SNPs in genes *CG43902* (Fig. 4C) and *ACC* (Fig. 4D), plasticity was different between new genotypes with minor versus major allele. We did not validate the effect of candidate gene *CG43117* on abdomen size plasticity (Fig. S7G, Table 2, Dataset 2). For all the confirmed candidates for plasticity, the genotypes with impaired gene function were more plastic than their respective controls, and the DGRP genotypes harboring the minor allele were more plastic than those harboring the major allele (Fig. S7). We also validated three out of four candidate QTLs for within-temperature variation in body size (Fig. 4 and S7, Table 2, Dataset 2). Genes *Nmdmc* (Fig. 4G) and *Optix* (Fig. 4I) affected thorax size at 28°C and at 17°C, respectively, while a SNP in gene *CG14688* (Fig. 4H) affected thorax size at 28°C. We did not validate the effect of candidate gene *Wapl* on abdomen size at 28°C (Fig. S7K, Table 2, Dataset 2).

To explore the pleiotropic effect of validated plasticity QTLs, we investigated whether the plastic response was also seen in the body part for which the SNP/gene had not been significantly associated to in the GWAS analysis (Fig. S7, Table 2). Out of the six validated plasticity QTLs, we only found cross-body part effects for gene *btv;* initially implicated in variation in plasticity of thorax size, this gene was also found to influence variation in plasticity of abdomen size (Fig. S7B, Table 1).

## DISCUSSION

Body size and body size proportions are traits closely associated to fitness (1, 2, 5), which vary greatly between species and populations, as well as between sexes and between same-sex individuals (11, 39, 61). Environmental conditions, such as temperature or food availability, can work both as inter-generational selective agents impacting body size evolution, and as intra-generational instructive agents that affect body size during development (7, 9, 33, 62). Studies in various species have explored what shapes inter- and intra-specific differences in size, including the physiological basis of body size regulation (16, 36, 38, 63) and the genetic basis of variation in body size (6, 64, 65).

### Components of variance in body size

Studying a panel of populations representing naturally-segregating alleles, the DGRP, we quantified effects of G, E, and GxE interactions on the size of two body parts (thorax and abdomen). As is well documented for various species of insects (32, 66, 67), most *D. melanogaster* genotypes we analyzed showed larger bodies when flies were reared at our lower temperature. However, we also documented cases of genotypes showing no plasticity (robustness) or showing size plasticity in the opposite direction. We also found a positive correlation in the levels of plasticity for the two body parts. The strong associations between the sizes of different body parts and plasticity therein is likely to reflect the tight regulation of body proportions, which is key for organismal performance (7, 68).

It is unclear to what extent a genotype’s responsiveness to environmental conditions (i.e. its plasticity) is associated with inter-individual differences found within a given environment for that same genotype (quantified with the coefficient of variation). While the latter is presumably un-accountable for by the effects of G, E, or GxE, it could reflect small genetic differences between individuals (e.g. derived from somatic mutation), micro-environmental variation (e.g. differences within a vial), or stochasticity in phenotype expression (e.g. developmental noise). We found that genotypes that were more plastic for thorax size (but not abdomen) also had higher levels of intra-genotype, intra-environment variation (i.e. higher coefficient of variation). Whether this component of phenotypic variance is assignable to micro-environmental variation and whether it has its own genetic basis has started to be investigated (53, 69, 70) and will, undoubtedly, be a topic of targeted future research.

### QTLs for size and size plasticity of different body parts

By using the raw and absolute values of the slopes of reaction norms as quantitative traits, we identified loci associated with variation in size plasticity. Genetic variation for environmental responsiveness could, in principle, involve different types of molecular players and could affect multiple traits in a similar or different manner. We described QTLs influencing size plasticity corresponding to different functions in terms of putative SNP effects (e.g. missense, regulatory, or synonymous mutations; Tables 2 and S2), and described molecular function and biological process for corresponding genes. These genes could potentially be mediating environmental effects at different levels, from the perception of the environmental cue (e.g. gene *btv*, which has been implicated in sensory perception (71)), to the transmission of that information to developing tissues (e.g. gene *Eip75B*, coding for an ecdysone receptor (72)), or the execution of the information on those tissues (e.g. genes *Men, ACC*, and *Hsp60A*, coding for two metabolic enzymes and a chaperone (72), respectively).

Presumably, genes higher up in the process of responding to the environment (e.g. those involved in the perception of external conditions versus those responding in specific tissues) would be more likely to affect multiple plastic traits in a concerted manner. With the exception of *btv*, none of our validated plasticity QTLs affected plasticity for other than the body part they had been identified as QTL for. Even for the complete set of candidate loci, we found very little overlap between QTLs for plasticity of different body parts (thorax versus abdomen), as well as for different properties of the reaction norms (raw versus absolute value of slopes). Furthermore, we also documented mostly private QTLs influencing variation in size at any given environment (i.e. body part and temperature-specific). Sex-, body part-, and environment-specific QTL effects had been previously documented for various traits in different models (51, 73–75). In *D. melanogaster*, for example, different loci have been associated to variation in bristle number in different body parts (75), and to variation in size in different environments (76). Such private QTLs can potentially facilitate independent evolution of the traits.

Previous works exploring the genetic basis of environmentally sensitive variation have mostly focused on investigating QTLs whose effect vary across environments (QTL-by-environment interactions) for a variety of traits in different species (65, 77–79). Much less attention has been paid to unraveling the allelic variants contributing to differences in plasticity itself. Exceptions include mapping of the genetic basis of thermal plasticity in life-history traits in *Caenorhabditis elegans* (80), of photoperiodic plasticity in multiple traits in *Arabidopsis thaliana* (81) and, more recently, of thermal plasticity in cold tolerance in *D. melanogaster* (53). Our results revealed little overlap between the QTLs that contribute to variation in trait within environments and the QTLs that contribute to variation in trait plasticity, assessed from the slope of reaction norms. We documented loci underlying variation in size plasticity (i.e. properties of reaction norms) that are different from those underlying variation in size at any temperature (i.e. at 17°C and at 28°C). These results shed light onto a long-standing discussion about the genetic underpinnings of plasticity, which argue that either the genetic control of phenotypic plasticity happens via specific loci determining plastic responses or via the same loci that control trait values at a given environment (82–85). Our data show that the genetic basis for trait plasticity, to a large extent, differs from the genetic basis for phenotypic variation in the trait itself.

### Evolution of size and size plasticity

Plasticity can be adaptive in that it helps populations cope with environmental heterogeneity, and it has even been argued that it can promote phenotypic and taxonomic diversification (20–22, 27). Theoretical models highlight the ecological conditions that should influence the evolution of plasticity, such as the predictability of environmental fluctuations (86) and costs of plasticity (87, 88). Plasticity is generally presumed to be costly and only selected for in predictably heterogeneous environments, such as seasons (89). The absence of a correlation between our thermal plasticity measurements and various fitness-related traits measured for the same genotypes could not identify any such cost. These potential costs might involve traits that have not been considered here, or these same traits but under (environmental) conditions that were not those assayed.

The ability to respond or resist environmental perturbation, and the balance between both processes, can be crucial for fitness in variable environments. In the DGRP, even though some degree of environmental responsiveness is maintained, we found that the alleles contributing to increased levels of plasticity occur nearly always at lower frequencies (i.e. the genotypes with the minor allelic variant having steeper reaction norms than those with the major allele). It is unclear to what extent this is the result of natural selection by the thermal regime that the natural population from which the DGRP was derived was exposed to, and/or the result of the process of deriving the mapping panel in the laboratory. It is also unclear to what extent QTLs for size plasticity in the DGRPs are those under selection in other populations. While it is conceivable, if not likely, that different QTLs contribute to variation in plasticity (or other quantitative traits) in different populations, we did find that a number of our plasticity QTLs have been targets of selection in other populations. Specifically, some of our candidate genes for size plasticity appear to have been selected in experimental populations of *D. melanogaster* evolving under different fluctuating thermal regimes (90). Among our 192 candidate QTLs for thermal plasticity, eight genes (including the validated *btv*) had changes in the populations evolving under hot and cold temperatures fluctuations (84), nine genes (including *Men*) had changes in populations evolving under hot fluctuations, and 25 genes had changes in the populations evolving under cold temperatures fluctuations (see all overlaps in Table S2). Gene *Eip75B* has also been previously implicated in the response to artificial selection for body size (91) and in differentiation between clinal populations (92, 93), which typically represent different thermal environments.

Altogether, our results shed light onto the nature of inter-genotype variation in plasticity, necessary for the evolution of plasticity under heterogeneous environments. We showed that QTLs for size plasticity: 1) bear alleles for increased plasticity at low frequencies, 2) correspond to polymorphisms in different genomic regions and within genes of a multitude of functional classes, and 3) are mostly “private QTLs”, with little overlap between our various GWAS analysis. The latter underscores the potential for independent evolution of trait and trait plasticity (different QTLs for size plasticity and for within-environment size variation), plasticity of different body parts (different QTLs for size plasticity of thorax and of abdomen), and even properties of the environmental response (different QTLs for raw and absolute slopes of thermal reaction norms).

## MATERIALS AND METHODS

### Fly stocks and rearing conditions

Data for the GWAS was collected from adult female flies of the Drosophila Genetic Reference Panel (DGRP) obtained from the Bloomington Stock Center. The DGRP is a set of fully sequenced inbred lines collected from a single population in Raleigh, NC, USA (46, 48). The number and the details of the lines included in the GWAS for each trait can be found in Table S1. Mutant stocks for the functional validations were: *Hsp60A* (stock 4689 from Bloomington), *MenEx3* and *MenEx55* (obtained from the T. Merritt lab). Control genetic backgrounds were *w1118* (stock 5905, from Bloomington) and Canton-S (obtained from C. Mirth lab). *UAS-Gal4* and UAS-RNAi lines used for validations were: stocks 6803 for *bab-Gal4*, 5138 for *tub*-Gal4, 28737 for *btv-*RNAi, and 35785 for mCherry-RNAi (all obtained from the Bloomington stock center), and stocks v108399 and v110813 for *Eip75B* and *Optix-RNAi*, respectively (obtained from the VDRC stock center).

Fly stocks were maintained in molasses food (45 gr. molasses, 75gr sugar, 70gr cornmeal, 20 gr. Yeast extract, 10 gr. Agar, 1100 ml water and 25 ml of Niapagin 10%) in incubators at 25°C, 12:12 light cycles and 65% humidity until used in this study. For the experiments, we performed over-night egg laying from ~20 females of each stock in vials with *ad libitum* molasses food. Eggs were then placed at either 17°C or 28°C throughout development. We controlled population density by keeping between 20 and 40 eggs per vial. We reared 200 DGRP lines and quantified thorax and abdomen size of 5 to 20 females per line, temperature and replicate. For 129 DGRP lines, we ran two replicates and for 32 lines we ran three replicates. The total number of flies used varied among DGRP lines due to differences in mortality at one or both of the temperatures. For some specimens, we could only quantify size of one body part if, for example, the individual was not properly positioned in the image or was damaged. Full details on the stocks used and the number of flies used per stock and temperature can be found in Dataset 1 and Table S1. Rearing conditions for the validations of candidate QTLs were similar to those used for the DGRP lines.

### Phenotyping body size and plasticity

Adult female flies (8-10 days after eclosion) were placed in 2ml Eppendorf and killed in liquid nitrogen followed by manual shaking to remove wings, legs and bristles. Bodies were mounted on Petri dishes with 3% Agarose, dorsal side up, and covered with water to avoid light reflections. Images containing 10 to 20 flies were collected with a LeicaDMLB2 stereoscope and a Nikon E400 color camera under controlled imaging conditions of light and white-balance. Images were later processed with a customized Mathematica macro to extract size measurements. For this purpose, we drew two transects per fly, one in the thorax and one in the abdomen, using body landmarks (as shown in Fig. S1A). Size of each body part was initially quantified as the number of pixels in the transect and later converted to millimeters. For abdominal transects, when necessary, we performed an additional step that involved the removal of pixels corresponding to the membranous tissue that is sometimes visible between segments.

### Genome-Wide Association Study

For each body part (thorax and abdomen), we performed four independent Genome-Wide Analyses (GWAS): two for thermal plasticity (raw and absolute values of the slopes of the reaction norms), and two for within-environment variation (length at 17°C and length at 28°C). Slopes of the reaction norm were calculated as the slope of the regression model lm (*Size ~ Temperature*) for each body part and DGRP line. The GWAS for variation in thermal responsiveness were done by using the raw and absolute values of the reaction norms, testing the model lm (*Slope ~ Allele + (1|Wolb|DGRP)*), *Wolb* being the *Wolbachia* status of the DGRP lines (47, 45). The GWAS analyses for within-environment variation (at either 17°C or 28°C) were done by testing the model lm (*Size ~ Allele + (1|Wolb|DGRP)*). All the GWAS were performed by using SNPs where we had information for at least ten DGRP lines per allele. We did not find an effect of *Wolbachia* in any of our GWAS analyses.

For each of the GWAS, we annotated the SNPs with a p-value < 10e-5 using the FlyBase annotation (FlyBase release FB2017_05; (72)). For the same SNPs, we performed first, a gene ontology enrichment analysis using the publicly available GOrilla Software (94, 95) and second, a network enrichment analysis using gene-enrichment and pathway-enrichment analyses were done using the publicly available NetworkAnalyst Software (96, 97); using all nodes from first order network generated with IrefIndex Interactome settings.

We tested for the effect of the chromosomal inversions (In_3R_K, In_3R_P, In_2L_t, In_2R_NS and In_3R_Mo) on our thorax and abdomen traits by using the models lm (*Mean Size ~ Inversion)* for within-environment size variation and lm (*Slope ~ Inversion)* for size plasticity variation.

Genetic distance matrix for the DGRPs was obtained from http://dgrp2.gnets.ncsu.edu/data.html and was used to perform a cluster hierarchical dendogram using *ape* and *phylobase* R packages. We estimated the phylogenetic signal and statistical significance for each of our traits using Blomberg’s K (92) and Pagel’s λ (93) metrics with the *phylosig* function in the *phytools* R package (94).

### Functional validations

The subsample of significant QTLs to be validated was taken from a first list of candidates (Fig. S3 and S4) selected based on p-value and corresponding peaks in the Manhattan plots (clear peaks prioritized), putative effect (missense and regulatory variants prioritized over intergenic variants), associated genes (annotated and known function prioritized). We used three methods for validation, depending on QTL properties: null mutants and RNAi (Gal4/UAS system) for genes containing several significant SNPs and/or containing SNPs corresponding to missense variants, and Mendelian randomization (MR) for SNPs in genes with little or no information available. Mutant and RNAi test that no or low levels of peptide affect variation in the quantitative trait for which the gene was identified as a candidate QTL while MR tests for sufficiency and independence from genetic background of the specific allele. Following these criteria we tested a total of 11 candidate SNPs/genes.

Validations by null mutants were done by comparing the phenotype in the heterozygous mutant stock with its respective genetic background. Validations by RNAi were done by comparing, for each Gal4 driver line, the phenotype of the gene of interest knockdown with the corresponding control cross using UAS-mCherryRNAi as well as with the corresponding control genetic background for the UAS line. We always used two different driver lines for our validations by RNAi: tub-Gal4 and bab-Gal4. However, for all candidate genes selected for RNAi validation, except *Nmdmc*, the crosses between RNAi line and tub-Gal4 were lethal.

The identity of the SNPs tested by MR is given by their annotation with Genome Release v6. For each candidate SNP, we first selected 10 DGRP lines to make a population with the minor allele fixed and 10 others to make a population with the major allele fixed. The 10 DGRP lines used to create each population were checked for having only one of the significant QTLs (p<10-5) fixed and not the others. These lines were used to generate four populations, two fixed for the major allele and two for the minor allele (Table S1). Each population was established by crossing 8 virgin females from each of 5 of the same-allele lines to 8 males of the other 5 lines. Reciprocal crosses were used to set two independent populations per allele. These populations were allowed to cross for eight generations to randomize genetic backgrounds. We confirmed by Sanger sequencing that those populations had our candidate allele fixed. Primer sequences used to confirm the allele in each population were:

Gene *CG43902* - forward primer: ACCACCAACATCAGCGTTTC; reverse primer: TGGTTTCGGCGTAGTTGTTG.

Gene *ACC* – forward primer: TGGGAAAAACCGGCCTAAGA; reverse primer: ATTTGTGGCTGTGGATTGCG.

Gene *CG43117* – forward primer: TAAGCAAAATGTGGCGTGCA; reverse primer: TTAACATGGATCCTGCGCAC

Gene *CG14688* – forward primer: CATACTTTGACAGACGGCCG; reverse primer: CGGCTACATTGTCATCGAGG

### Statistical analyses

All statistical analyses were performed with R Statistical Package version 3.3.1 (95). We used a linear model to test for the effect of replicate (model lm (*Size ~ Body_part*Temperature*DGRP*Replicate*)), that was found to be non-significant (p-value > 0.05). For each body part, we used linear models to test for the effect of genotype (model lm (*Size ~ DGRP*)) or the interaction between genotype and temperature (model lm (*Size ~ DGRP*Temperature*)) on size. Reaction norms for each DGRP line were calculated by using the regression model lm (*Size ~ Temperature*). Using the results from that model, we defined plastic genotypes as those DGRP lines whose reaction norm slope was significantly different from zero (p-value < 0.05) and we extracted two properties of the reaction norms per DGRP line and body part: the absolute value of the slope as a measurement of thermal sensitivity, describing only the magnitude of the response to temperature, and the raw value of the slope as a measurement which describes also the direction of that response. Linear mixed models were calculated using *lme4* R package.

We used Pearson correlations (α = 0.99) to test for linear correlation in size between body parts, controlling for DGRP lines. We also used Pearson correlations to test for linear correlations among our measured traits and between those and other available datasets for the DGRPs. For this, we used the mean value per DGRP line for each trait and the *corrplot* R package. We report both correlation coefficient and significance levels. Available DGRP phenotypes that were used to correlate with our traits were: size measurements at 25°C (49), longevity (43), starvation resistance, chill coma recovery (45), tolerance to infection with *Providencia rettgeri* bacteria (54) and resistance to infection with *Metarhizium anisopliae* fungi or with *Pseudomonas aeruginosa* bacteria (48).

Broad sense heritability for size at each temperature was estimated as[inline]where[inline]and[inline] are the among-line and within-line variance components, respectively. Heritability of plasticity was calculated as[inline]where[inline]and[inline] are the variance associated with the genotype by environment interaction and total variance components, respectively, as proposed in Scheider and Lyman (1989). Variance components were extracted using *varcomp* R package.

For the functional validations of within-environment SNPs and genes we tested the model lm (*Size ~ Allele)* and lm (*Size ~ Genotype)*, respectively. For the validations of plasticity SNPs and genes we tested the model we tested the model lm (*Size ~ Genotype*Temperature)* and lm (*Size ~ Allele*Temperature)*, respectively. In all cases, significant differences among groups were estimated by post hoc comparisons (Tukey’s honest significant differences).

## DATA ACCESIBILITY

Raw data (Dataset 1 and 2) will be archived in the Dryad Digital Repository upon acceptance of the manuscript.

## AUTHORS’ CONTRIBUTIONS

E.L. and P.B. designed the study. E.L. collected the data. E.L. and D.D. analyzed the data. E.L. and P.B. wrote the manuscript, with contributions from D.D.

## ACKNOWLEDGEMENTS

We are grateful to Thomas Merrit for sharing the *Men* mutants, and to Daniel Sobral and IGC’s Bioinformatics Unit and Fly Unit for technical support. We also thank Erik van Bergen for comments on the manuscript. Computations were performed on the EDB-Calc Cluster, which uses a software developed by the Rocks(r) Cluster Group (San Diego Supercomputer Center, University of California, San Diego and its contributors), and is hosted by the laboratory “Evolution et Diversité Biologique” (EDB) and supported by Pierre Solbes.

